# Aging is associated with a modality-specific decline in taste

**DOI:** 10.1101/2024.02.01.578408

**Authors:** Elizabeth B. Brown, Evan Lloyd, Alfonso Martin-Peña, Samuel McFarlane, Anupama Dahanukar, Alex C. Keene

## Abstract

Deficits in chemosensory processing are associated with healthy aging, as well as numerous neurodegenerative disorders, including Alzheimer’s Disease (AD). In many cases, chemosensory deficits are harbingers of neurodegenerative disease, and understanding the mechanistic basis for these changes may provide insight into the fundamental dysfunction associated with aging and neurodegeneration. The fruit fly, *Drosophila melanogaster*, is a powerful model for studying chemosensation, aging, and aging-related pathologies, yet the effects of aging and neurodegeneration on chemosensation remain largely unexplored in this model, particularly with respect to taste. To determine whether the effects of aging on taste are conserved in flies, we compared the response of flies to different appetitive tastants. Aging impaired response to sugars, but not medium-chain fatty acids that are sensed by a shared population of neurons, revealing modality-specific deficits in taste. Selective expression of the human amyloid beta (Aβ) 1-42 peptide bearing the Arctic mutation (E693E) associated with early onset AD in the neurons that sense sugars and fatty acids phenocopies the effects of aging, suggesting that the age-related decline in response is localized to gustatory neurons. Functional imaging of gustatory axon terminals revealed reduced response to sugar, but not fatty acids. Axonal innervation of the fly taste center was largely intact in aged flies, suggesting that reduced sucrose response does not derive from neurodegeneration. Conversely, expression of the amyloid peptide in sweet-sensing taste neurons resulted in reduced innervation of the primary fly taste center. A comparison of transcript expression within the sugar-sensing taste neurons revealed age-related changes in 66 genes, including a reduction in odorant-binding protein class genes that are also expressed in taste sensilla. Together, these findings suggest that deficits in taste detection may result from signaling pathway-specific changes, while different mechanisms underly taste deficits in aged and AD model flies. Overall, this work provides a model to examine cellular deficits in neural function associated with aging and AD.

## INTRODUCTION

Aging is associated with deficits in numerous sensory modalities including taste and smell [1–4]. In humans, several neurodegenerative diseases including Alzheimer’s Disease (AD) and Parkinson’s Disease (PD) have been shown to disrupt primary sensory cells, though less is known about the physiological and functional effects of healthy aging[3,5]. Most neurodegenerative diseases exhibit chemosensory deficits that precede motor or memory deficits, which are hallmarks of PD and AD respectively [6,7]. The taste system provides a particularly tractable system for examining the effects of aging on chemosensory processing. Unlike olfaction, taste is comprised of relatively few modalities that elicit reflexive responses. Investigating how taste response changes in aging and AD model animals has the potential to identify the genetic and neural basis contributing to age-related chemosensory decline.

The fruit fly, *Drosophila melanogaster*, has developed into a powerful model to study taste processing because of its amenability to genetic manipulation and functional parallels in taste processing between flies and mammals [8–10]. Additionally, the gustatory system of *Drosophila* is amenable to *in vivo* Ca^2+^ imaging and electrophysiology, both of which can be coupled with robust behavioral assays that measure reflexive taste response and food consumption [11]. Among the main classes of non-overlapping gustatory neurons are sweet-sensing neurons, which promote feeding and are labeled by expression of the *Gustatory Receptor 64f (Gr64f)*; these neurons respond to both sugars and fatty acids [12–14]. The well-defined function and projection of these neurons provide a system to examine how gustatory function changes in normal and pathological aging.

The short lifespan of fruit flies provides a system for examining the genetic basis of age-related deficits in many behavioral and physiological traits. The effects of aging are accelerated in the neurodegenerative disease model *Drosophila* expressing amyloidogenic forms of human Amyloid β (Aβ) and recapitulate several key features of AD, including Aβ accumulation, age-dependent learning impairment, and neurodegeneration [15–17]. In a commonly used model, a variant of human Aβ_1-42_ harboring the mutation E22G, termed the Arctic mutation, induces robust neurodegeneration as well as behavioral deficits [17–21]. While most experiments examining the effects of AD variants in flies drive AD-associated variants throughout the brain, targeted expression of disease-causing variants in defined populations of neurons allows for investigation of how aging and Aβ expression impact the connectivity and physiology of individual circuits.

Here, we examine the effects of aging or Arctic-bearing Aβ_1-42_ expression on gustatory function and behavior. We find that both healthy aging and pathological aging through cell-autonomous expression of Arctic in sweet taste neurons result in deficits in the detection of sugars, but not fatty acids. These tastants are detected by shared neurons but are activated by unique taste receptors and intercellular signaling pathways. Our findings suggest that both aging and Arctic-bearing Aβ_1-42_ expression disrupt cellular physiology in a modality-specific manner. We also identify many candidate regulators of age-related deficits in taste. Therefore, gustatory neurons provide a model to study the mechanisms underlying aging and the neurodegeneration-related decline of sensory processing.

## RESULTS

To examine the relationship between aging and taste, we measured the proboscis extension response (PER) against varying concentrations of sucrose in young *w*^1118^ 10-day-old flies and aged 40-day-old flies. In this assay, a tastant is provided to the proboscis, eliciting a reflexive response independent of post-ingestive feedback (Fig 1A; [22]). To define response sensitivity, flies were provided with different concentrations of sucrose ranging from 1mM to 1M. In 10-day-old flies, the response to these concentrations ranged from 15% to 100%, providing a range to measure taste sensitivity. The sucrose response did not differ at concentrations of 1mM and 1M between young and aged flies; however, there was a significant decrease in responsiveness to sucrose in aged flies at intermediate concentrations of 20mM and 50mM (Fig 1B). To determine whether this aging-dependent decline in taste sensitivity to sucrose can be generalized to other sweet tastants, we measured PER to different concentrations of fructose. Like our results using sucrose, we found that taste sensitivity to fructose significantly declines at intermediate concentrations (10mM and 20mM; Fig S1A). The finding that sugar taste does not differ at high concentrations suggests that flies are capable of recognizing sugar and performing the motor task of full proboscis extension, even though sensitivity to sweet tastants is diminished in aged animals.

**Figure 1.**
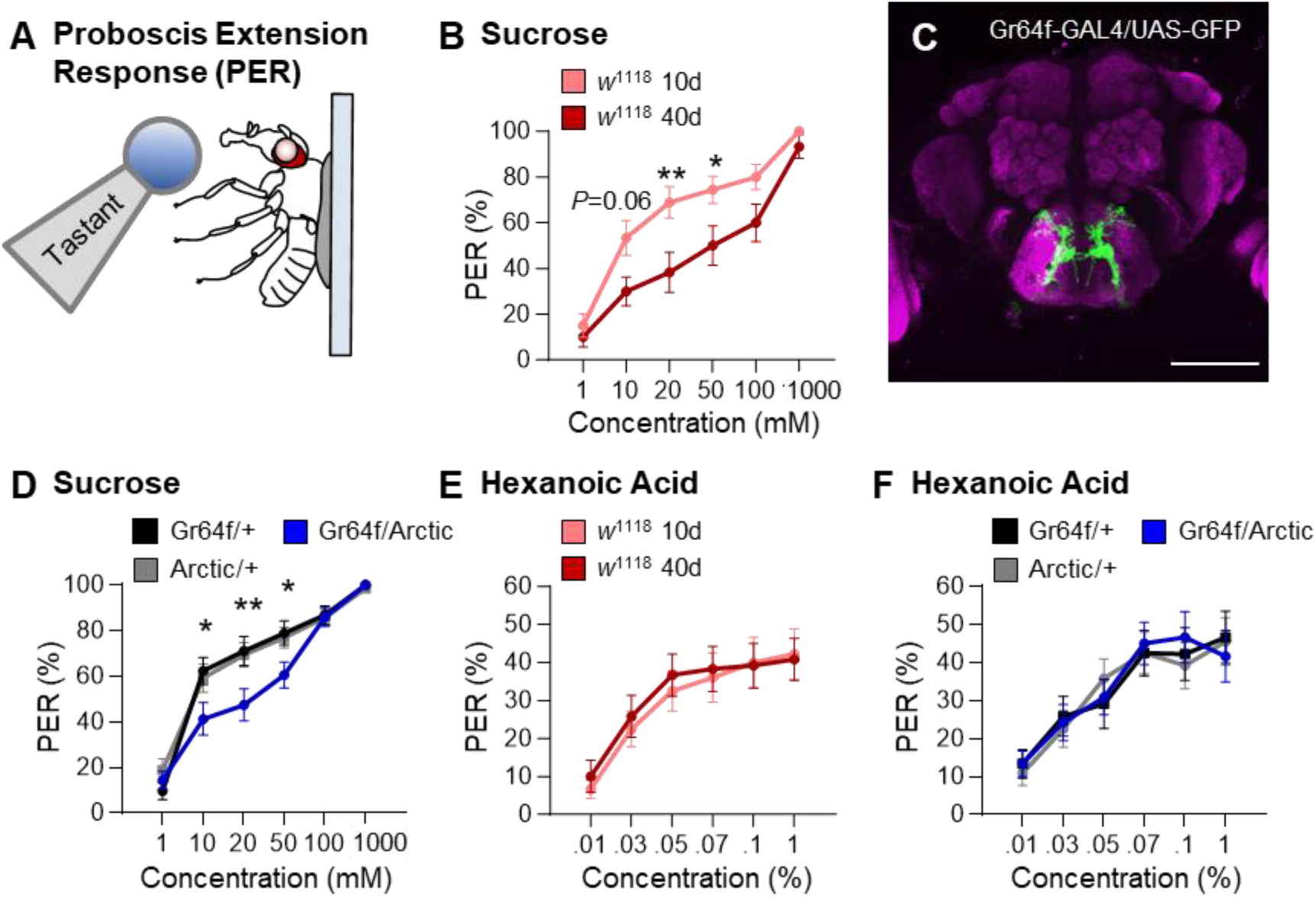
Taste response to sugar, but not fatty acids, is reduced at intermediate concentrations in aging and AD model flies. **(A)** Proboscis extension response (PER) was measured in female flies after 24 h of starvation. Either sucrose or hexanoic acid was applied to the fly’s labellum for a maximum of 2 s and then removed to observe proboscis extension reflex. **(B)** There is a significant effect of age on PER to sucrose in *w*^1118^ flies (two-way ANOVA: F_1,278_ = 23.77, *P* < 0.0001; N = 20–30). **(C)** Expression pattern of *Gr64f* is visualized with GFP. *Gr64f*-expressing neurons project to the subesophageal zone of the brain. Background staining is NC82 antibody (magenta). Scale bar = 100 μm. **(D)** There is a significant effect of Arctic expression on PER to sucrose (two-way ANOVA: F_2,702_ = 7.529, *P* < 0.0006; N = 28–49). **(E)** There is no effect of age on PER to hexanoic acid in *w*1118 flies (two-way ANOVA: F_1,415_ = 0.2959, *P* < 0.5867; N = 20–40). **(F)** There is no effect of Arctic expression on PER to hexanoic acid (two-way ANOVA: F_2,671_ = 0.0396, *P* < 0.9612; N = 30–40). Error bars indicate ± SEM. ^*^*P* < 0.05; ^**^ *P* < 0.01.

In *Drosophila*, PER involves the activation of a sensory circuit that includes gustatory neurons, interneurons, motor neurons, and musculature [11,23,24]. In flies, broad expression of the amyloid β (Aβ_1-42_) variant Arctic recapitulates many aspects of premature aging including disrupted sleep, memory impairment, and premature death [17–21,25,26]. To determine whether dysfunction in the sensory component of the taste circuit alone is sufficient to drive this decline in taste sensitivity, we expressed Arctic exclusively in sweet taste neurons labeled by the *Gustatory receptor 64f* (*Gr64f*; Fig 1C). We tested 20-day-old flies because this is prior to age-related loss of taste in controls, allowing for the identification of accelerated loss of taste. PER was significantly diminished in 20-day-old flies expressing Arctic in sweet taste neurons (*Gr64f*-GAL4>UAS-Arctic) compared to controls harboring *Gr64f*-GAL4 or UAS-Arctic alone (Fig 1D). Similarly, a decline in taste sensitivity in flies that express Arctic in *Gr64f* neurons was also observed in response to fructose (Fig S1B). Further, there was no effect on the expression of the non-toxic 40-amino acid form of Aβ (Aβ_1-40_) to sucrose presentation (Fig S1C), suggesting that the results observed with Arctic are related to the pathogenicity of Aβ_1-42_ [27,28]. Therefore, Arctic expression induces cell-autonomous deficits in sweet taste response, phenocopying the effects of aging and supporting the notion that reduced sensitivity of taste neurons underlies aging-related reductions in PER.

The *Gr64f*-expressing taste neurons detect numerous appetitive tastants, including sugars and fatty acids [29–31]. While these neurons are activated by both sugars and fatty acids, the response to tastants is conferred by distinct signaling pathways with sugar response dependent on the Gsα subunit [32,33], while fatty acid response is dependent on parallel signaling pathways that include Phospholipase C and *Ionotropic Receptor 56d* (*Ir56d*; [29,30,34]). To determine whether aging impairs response to both tastants, we compared the response of young and aged flies to hexanoic acid, a medium-chain fatty acid that robustly activates *Gr64f* neurons (Fig 1E; [30,31]. Across five different concentrations ranging from 0.01% to 1%, there was no difference in response to hexanoic acid. Higher concentrations of this tastant are aversive; therefore, they were not tested in this assay [29]. Together, these findings suggest that fatty acid taste is not impaired in aged animals.

To determine whether Arctic expression in *Gr64f* neurons impairs fatty acid taste, we measured taste response to hexanoic acid in 20-day-old flies expressing UAS-Arctic in taste neurons (Fig 1F). There was no difference in PER between control flies harboring *Gr64f* and experimental flies expressing Arctic in *Gr64f* neurons (*Gr64f*-GAL4>UAS-Arctic). We also measured PER to different concentrations of octanoic acid, a second fatty acid that is also activated by *Gr64f* neurons [31]. There was no significant difference in taste response to octanoic acid between young and aged flies expressing Arctic in *Gr64f* neurons (Figure S1D, E). Similarly, expression of the non-toxic Aβ_1-40_ did not affect taste response to hexanoic acid (Fig S1F). Therefore, both natural aging and taste neuron-specific expression of Arctic impairs response to sugars without affecting fatty acid taste response.

We next sought to determine whether the observed differences in aging and AD model flies are sex specific. We found that the sensitivity to sucrose, but not hexanoic acid, is reduced in 40-day-old male flies compared to their 10-day-old counterparts (Fig S2A, B). Similarly, expression of Arctic in *Gr64f* neurons in 20-day-old male flies impairs sensitivity to sucrose, but not fatty acids (Fig S2C, D). Therefore, both aging and Arctic expression selectively impact sucrose response in male and female flies.

It is possible that the decline in taste response is due to neurodegeneration of the taste neurons, loss of connectivity with taste neuron targets, or reduced responsiveness of taste neurons. Given that the decline in taste response is modality-specific, we hypothesize that neural responsiveness to sugars, but not to and fatty acids change with age. To test this hypothesis, we first measured neural activity of *Gr64f* neurons to tastant presentation (Fig 2A). To directly assess the effects of aging on neural activity, we expressed the genetically encoded Ca^2+^ sensor GCaMP6s-mCherry (UAS-Gerry) in *Gr64f* neurons. Quantifying the ratio of stable mCherry signal to Ca^2+^ induced GCaMP6s provides a ratiometric quantification of evoked activity [35–37]. We applied an in vivo preparation to record responsiveness of projections in the subesophogeal zone (SEZ) when tastants were applied directly to the proboscis (Fig 2B; [30]). At concentrations ranging from 10mM to 100mM, the responses of 40-day-old flies to sucrose were significantly reduced compared to 10-day-old flies (Fig 2C.D). To determine if aging also impacts fatty acid taste response, we compared the response in the SEZ to 0.01%, 0.1%, and 1% hexanoic acid in 10-day-old and 40-day-old flies (Fig 2C). Across all three concentrations, there were no differences in Ca^2+^ response between 10- and 40-day-old flies (Fig 2D). Therefore, consistent with our behavioral results, aging diminishes taste neuron response to sucrose, but not hexanoic acid presentation.

**Figure 2.**
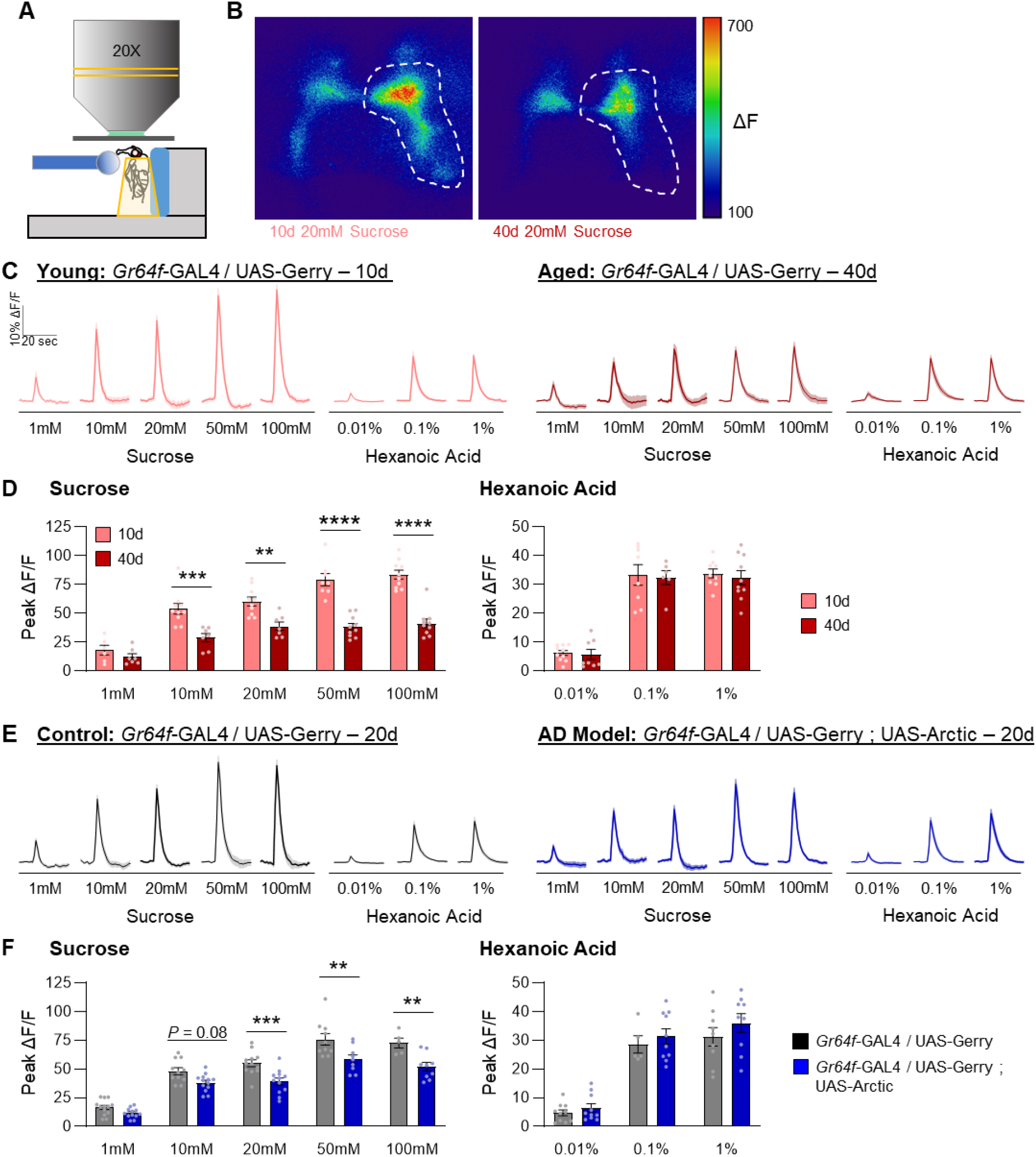
Neuronal activity in *Gr64f* neurons is reduced during aging and in an AD model upon sucrose but not hexanoic acid presentation. **(A)** Diagram of live-imaging experimental design. A tastant is applied to the proboscis while florescence is recorded simultaneously. **(B)** Representative pseudocolor images of GCaMP/mCherry fluorescence in *Gr64f* neurons in response to 20mM sucrose in 10 (left) and 40 day-old flies (right). The peak in UAS-GCaMP6 fluorescence is shown. Scale bar = 50 μm. **(C)** The average ratio of GCaMP/mCherry of *Gr64f* neurons in response to each tastant and concentration in 10 and 40 day old flies. The shaded region of each trace indicates ± SEM. **(D)** Average peak change in the ratio of GCaMP/mCherry for data shown in **(C)**. There is a significant effect of age on neuronal response to sucrose (two-way ANOVA: F_1,75_ = 112.1, *P* < 0.0001; N = 6– 10), but not to hexanoic acid (two-way ANOVA: F_1,45_ = 0.2846, *P* < 0.5963; N = 6–10). **(E)** The average ratio of GCaMP/mCherry of *Gr64f* neurons in response to each tastant and concentration in control and AD model flies. **(F)** Average peak change in the ratio of GCaMP/mCherry for data shown in **(E)**. There is a significant effect of Arctic expression on neuronal response to sucrose (two-way ANOVA: F_1,96_ = 47.80, *P* < 0.0001; N = 5–13), but not to hexanoic acid (two-way ANOVA: F_1,47_ = 2.218, *P* < 0.1431; N = 5–10). For traces, the shaded region indicates ± SEM. For bar graphs, error bars indicate ± SEM. ^**^*P* < 0.01; ^***^ *P* < 0.001; ^****^ *P* < 0.0001.

To determine if the effects of aging are shared in flies with cell-autonomous expression of Arctic in taste neurons, we expressed UAS-Gerry in taste neurons and measured neural activity to sucrose and hexanoic acid presentation in 20-day-old flies (Fig 2E). Consistent with the results comparing 10 and 40 day old flies, the response to sucrose was reduced at intermediate (20mM) and high (100mM) concentrations in experimental flies expressing Arctic (*Gr64f*>UAS-Arctic) compared to age-matched flies harboring *Gr64f*-GAL4 alone (Fig 2F). There were no significant differences in response to hexanoic acid at any of the concentrations tested (Fig 2F). Therefore, selective expression of Arctic in sweet-sensing taste neurons has a selective effect on lowering sugar sensitivity, similar to the effects of aging on taste response. Taken together, the reduced behavioral response to sucrose in both aged and Arctic flies may be attributed to the observed deficit in *Gr64f* neurons.

It is also possible that reductions in synaptic innervation of the suboesophageal zone contribute to age-dependent loss of sucrose responsiveness. To explore this possibility, we directly measured innervation of the SEZ *Gr64f* neurons. We expressed the membrane-bound marker mCD8GFP in *Gr64f* neurons and quantified immunofluorescence from sensory terminals that innervate the SEZ at days 10 and 40 (Fig 3A). The *Gr64f* neurons project to distinct regions of the SEZ, with neurons from the labellum projecting to the ventral region of the SEZ, while neurons from the taste pegs and internal mouthparts primarily project to the dorsal region of the SEZ [35– 37]. The volume and intensity within these two regions were increased in both labellum and taste peg regions (Fig 3B, C). This increase in intensity may be attributable to either the accumulation of membrane-bound GFP at the synapse during aging, or to a strengthening of connections. Together, these results suggest that the reduced PER in response to sucrose is not due to large-scale neurodegeneration or loss of SEZ innervation.

**Figure 3.**
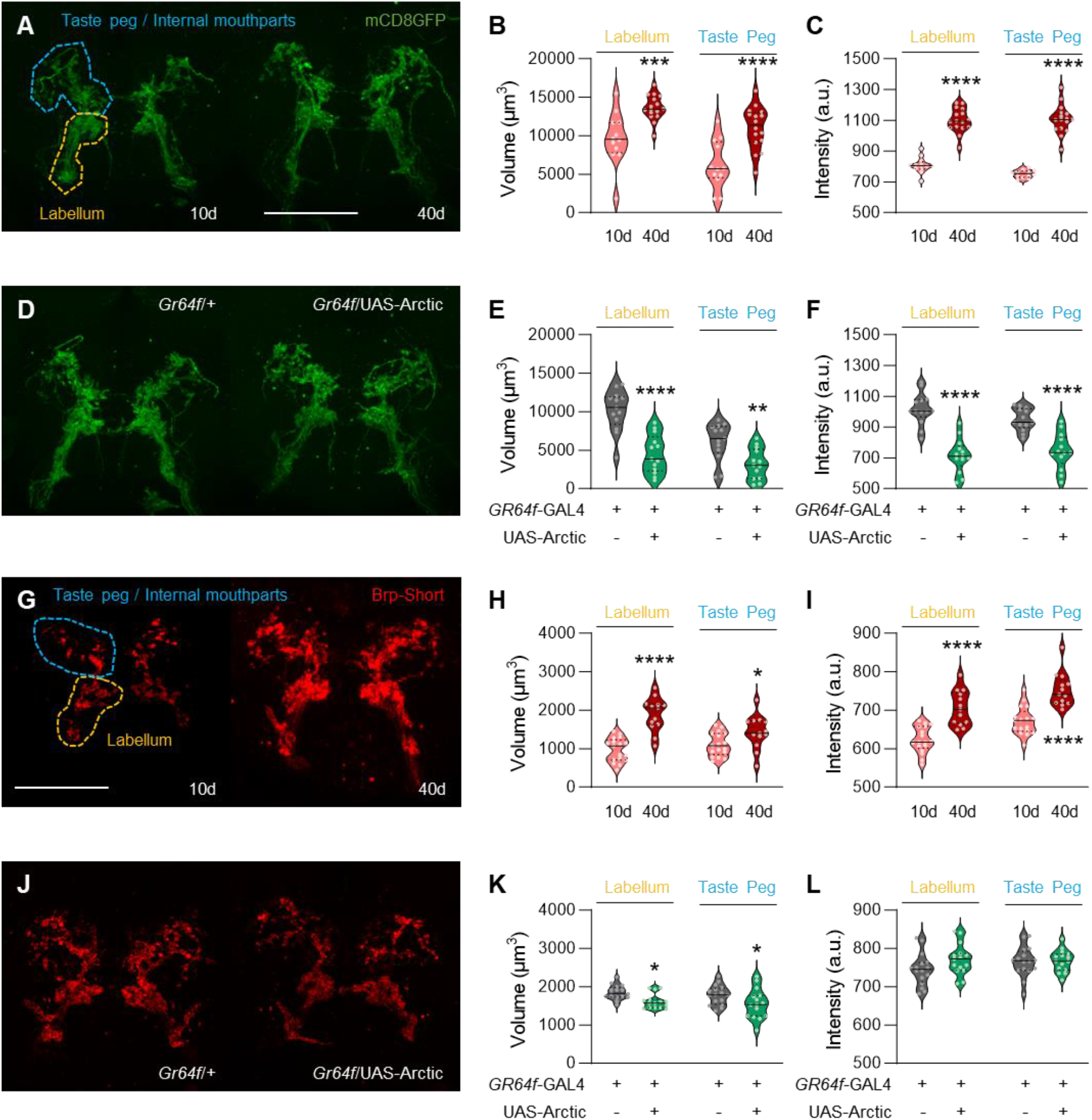
Aging and Arctic expression have differential effects on the axon terminals and active zone scaffold of *Gr64f* neurons. **(A)** Expression pattern of *Gr64f* in 10 and 40 day old flies is visualized with GFP (green). ROIs indicating the labellar (yellow) and the taste peg/internal mouthpart projections (blue) are shown. Scale bar = 50 μm. **(B)** There is a significant effect of age on *Gr64f* neurite volume (two-way repeated measures ANOVA: F_1,26_ = 23.29, *P* < 0.0001). **(C)** There is a significant effect of age on *Gr64f* neurite intensity (two-way repeated measures ANOVA: F_1,26_ = 167.5, *P* < 0.0001). **(D)** Expression pattern of *Gr64f* in control and AD model flies is visualized with GFP (green). **(E)** There is a significant effect of Arctic expression on *Gr64f* neurite volume (two-way repeated measures ANOVA: F_1,24_ = 22.98, *P* < 0.0001). **(F)** There is a significant effect of Arctic expression on *Gr64f* neurite intensity (two-way repeated measures ANOVA: F_1,24_ = 45.01, *P* < 0.0001). **(G)** The active zone of *Gr64f* neurons in young and aged flies is visualized with Brp-Short (red). ROIs indicating the labellar (yellow) and the taste peg/internal mouthpart projections (blue) are shown. Scale bar = 50 μm. **(H)** There is a significant effect of age on *Brp* volume (two-way repeated measures ANOVA: F_1,28_ = 26.18, *P* < 0.0001). **(I)** There is a significant effect of age on *Brp* intensity (two-way repeated measures ANOVA: F_1,28_ = 43.09, *P* < 0.0001). **(J)** The active zone of *Gr64f* neurons in control and AD model flies is visualized with Brp-Short (red). (K) There is a significant effect of Arctic expression on *Brp* volume (two-way repeated measures ANOVA: F_1,29_ = 8.244, *P* < 0.0076). **(L)** There is no effect of Arctic expression on *Brp* intensity (two-way repeated measures ANOVA: F_1,29_ = 1.706, *P* < 0.2018). The median (solid line) as well as 25th and 75th percentiles (dotted lines) are shown. a.u. = arbitrary units. ^*^ *P* < 0.05; ^**^*P* < 0.01; ^***^ *P* < 0.001; ^****^ *P* < 0.0001.

We also sought to directly compare the effects of cell-autonomous Arctic expression on innervation of *Gr64f* axon terminals. A comparison between *Gr64f* controls and flies expressing Arctic in *Gr64f* neurons (*Gr64f*-GAL4>UAS-Arctic) revealed a significant reduction in both volume and intensity in projections from the labellum as well as from the taste pegs and internal mouthparts of 20 day-old flies (Fig 3D-F). Therefore, reduced synaptic innervation of axon terminals that project from the labellum may contribute to the reduced response to sucrose in flies that express Arctic in *Gr64f* neurons. Together, these findings raise the possibility that different mechanisms underlie natural aging- and Arctic-dependent decline in taste response to sugar.

It is possible that the change in signal intensity of *Gr64f* neurons with age and Arctic expression is driven by changes in the active zone scaffold, as it has been previously shown that aging substantially increases levels of the active zone component *Bruchpilot* (*Brp*) [38–41]. To label the active zones of *Gr64f* neurons, we expressed UAS-*Brp*-short [42], a nonfunctional Brp that localizes to endogenous Brp sites. We quantified Brp at *Gr64f* terminals of 10- and 40-day-old flies (Fig 3G). Both the volume and intensity of synaptic Brp levels in the labellar and taste peg regions were significantly elevated in aged flies, suggesting that the loss of synaptic proteins does not account for the reduced response to sucrose (Fig 3H, I). To determine whether these aging-dependent changes to the SEZ are similar in AD model flies, we next compared the effects of cell-autonomous Arctic expression on Brp levels in *Gr64f* neurons (Fig 3J). We found that Arctic expression in *Gr64f* neurons (*Gr64f*-GAL4>UAS-Arctic, UAS-Brp-short) significantly decreases volume, but not signal intensity in 20 day-old flies (Fig 3K, L). These findings suggest that aging and cell-autonomous Arctic expression have differential effects on innervation of the SEZ and assembly of the active zone in *Gr64f* neurons, raising the possibility that different mechanisms regulate the decline in taste response to sugar in aging and AD model flies.

To identify genes associated with the aging of chemosensory neurons, we performed 10x genomics-based single-nucleus RNA sequencing (snRNA-seq) [43] on the labellar tissue of young (10 day) and aged (40 day) flies (Fig 4A). This approach allowed us to examine the transcriptomes of individual sweet taste neurons and identify changes in gene expression associated with aging. Two independent rounds of sequencing were performed for each young and aged condition (with ∼400 labellum per replicate) to obtain a median of 1095 reads per nucleus. After filtering and correcting for batch effects, there were a total of 36194 young nuclei and 42639 aged nuclei in the pooled replicates (Fig 4B, C). The pooled replicates from young nuclei have a median of 558 genes and 1116 Unique Molecular Identifiers (UMIs) per cell, while the pooled replicates from aged nuclei have a median of 557 genes and 1080 UMIs per cell. The detected numbers of expressed genes and UMIs were largely consistent between young and aged pooled replicates (Fig S3).

**Figure 4.**
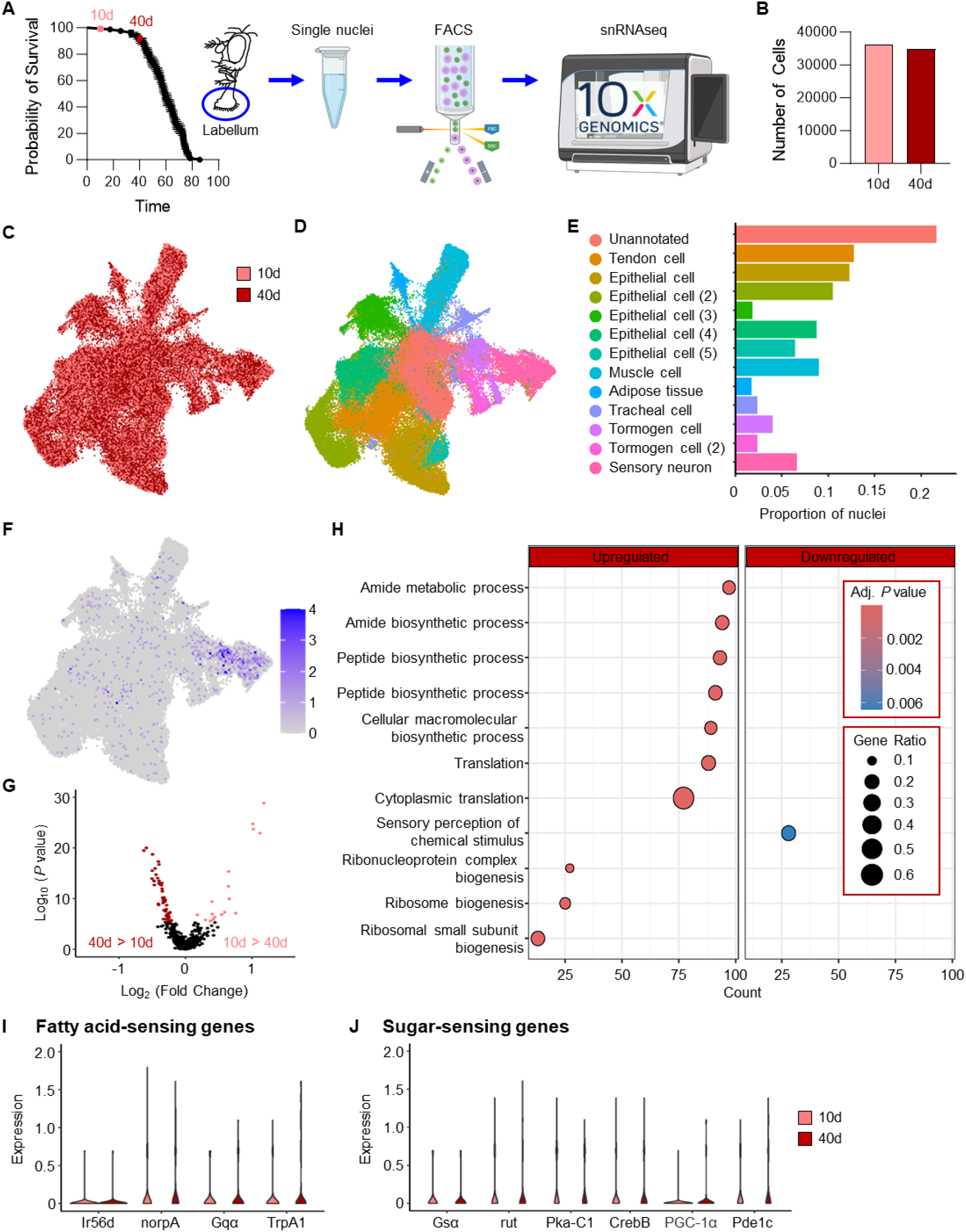
snRNA-sequencing of the labellum in young and aged flies. **(A)** Schematic of snRNA-sequencing workflow. The labellum of adult flies aged 10 or 40 days were dissected and dissociated to obtain single nuclei for 10x genomics snRNA-seq. Error bars indicate ± SEM. FACS: Florescence-activated cell sorting. **(B)** Number of nuclei collected from each age and replicate. **(C)** Uniform Manifold Approximation and Projection (UMAP) visualization of the 10- and 40-day-old, pooled replicate samples. **(D)** UMAP visualization representing unique clusters. Each color and dot on the plot represent a unique cluster and nucleus, respectively. **(E)** The proportion of nuclei for each broad cell cluster shown in **(D). (F)** UMAP visualization representing the average expression of *Gr64f* in 10d and 40d samples. **(G)** Volcano plot depicting differentially expressed genes within the *Gr64f*+ nuclei between 10- and 40-day-old flies. **(H)** Gene Ontology (GO) analysis of the differentially expressed genes identified in **(G). (I, J)** Gene expression of candidate genes associated with the **(I)** *Phospholipase C* signaling and **(J)** cAMP signaling pathways between the 10- and 40-day-old groups. There were no significant differences in gene expression.

To identify which types of nuclei are present among the pooled replicates, we performed clustering analysis and identified 13 distinct clusters, including a sensory neuron cluster (Fig 4D, E, bright pink). These clusters were identified based on known marker genes for each of these tissue types. In the case of sensory neurons, the top three marker genes include *paralytic* (*para*), *long non-coding RNA:noe* (*lncRNA:noe*), and *Shaker* (*Sh*; Fig S4A). Additional markers of this cluster include neural-specific genes, such as *Synaptobrevin* (*nSyb*), *Bruchpilot* (*Brp*), *Synaptotagmin 1* (*Syt1*), and *Cadherin-N* (*CadN*), as well as numerous odorant-binding proteins (OBPs) (Supplementary Table 1). We next sought to examine the effect of aging on gene expression within the sensory neuron cluster. To identify genes that change expression between the young and aged conditions, we performed differential gene expression analysis (Fig S4B, Supplementary Table 2). We identified 92 differentially expressed genes (DEGs), with 65 genes upregulated and 27 genes downregulated in the aged condition. We next assessed whether there was statistical enrichment of Gene Ontology (GO) terms among these DEGs (Fig S4C, Supplementary Table 3). We found significant upregulation of genes associated with translation and ribosome biogenesis, while significant downregulation was observed for genes involved in the perception of chemical stimuli. Overall, these findings suggest increased translation as a possible compensatory mechanism for reduced chemosensory expression.

Gene expression analysis of individual nuclei provides an opportunity to investigate the effect of aging on gene expression, specifically within *Gr64f*+ sweet taste neurons. We identified 404 and 360 nuclei that express *Gr64f* in the young and aged pooled replicates, respectively (Fig 4F). To identify transcriptomic differences between the young and aged Gr64f+ nuclei, we performed differential gene expression analysis and identified 66 DEGs with 49 genes upregulated and 17 genes downregulated in the aged condition (Fig 4G, Supplementary Table 4). There was substantial overlap between the *Gr64f*+ DEGs and those identified in the sensory cluster. We found that 63 of these genes overlap, suggesting that the effects of aging on sensory neurons may be generalizable to other taste neurons. These include the upregulated gene *Activity-regulated cytoskeleton associated protein 1* (*Arc1*) that has retrovirus function and is involved in transsynaptic communication [44], as well as multiple genes involved in the innate immune response, e.g., *Listericin* and *virus-induced RNA 1* [45–47]. We also identified upregulation of 38 ribosomal proteins. Further, numerous OBPs were found to be downregulated with age, raising the possibility that they are directly involved in diminished taste. We next assessed whether there was statistical enrichment of GO terms among these DEGs and found significant downregulation of genes involved in perception of chemical stimuli, while those associated with translation and ribosome biogenesis were significantly upregulated (Fig 4H, Supplementary Table 5). These findings further support the notion that changes in *Gr64f*+ nuclei may be generalizable to other sensory nuclei.

Different signaling pathways are responsive to fatty acids and sugars, with sugars activating the cAMP signaling pathway [32] and fatty acids activating the *Phospholipase C* signaling pathway [34]. To determine whether aging affects the expression of genes associated with either sugar or fatty acid taste perception, we examined genes whose expression has been previously implicated in either of these pathways (Fig 4I, J; [29–33,48,49]. However, there were no significant differences in expression observed in the examined genes with respect to either of these pathways. However, given the low detection rate of these genes, we cannot rule out the possibility that aging affects the expression of other aspects of these pathways.

## DISCUSSION

Our findings reveal that aging in *Drosophila* is associated with reduced sensitivity to sugar taste. There is an abundance of evidence that both olfactory and taste sensitivity is impaired during aging in humans and rodents [50–52]. In humans, the reduced sensitivity to tastants appears to occur across taste modalities, suggesting a generalized effect of normal aging on taste [4]. However, studies in mice reveal selective reductions in some modalities, but not others; for example, a reduction has been observed in the number of sweet taste receptor subunits. Taste sensitivity loss appears to be selective to sugar, but not other tastants including salt, acid, and bitter substances [52]. Therefore, our findings that age-related taste loss in *Drosophila* occurs in response to sugars but not fatty acids reveals that the effects of modality-specific taste loss in aged animals are present across both mammalian and non-mammalian species.

In humans, chemosensory deficits are present in many neurodegenerative diseases [53–55]. For example, a decline in olfactory function is reported to precede other neurological deficits in both AD and PD [56,57], suggesting that olfactory receptor neurons are highly sensitive to neurodegenerative processes. Additionally, taste perception is disrupted, and taste bud number is reported to be reduced in AD model mice, though other studies in different AD mice models have found no difference in taste [58,59]. However, studies in both humans and rodent models have not specifically examined whether the effects on taste are due to the primary sensory cells themselves, or to downstream factors. We expressed the AD E693G variant of Aβ, Arctic, selectively in taste neurons to determine whether cell-autonomous expression of this variant phenocopies the taste deficits observed during natural aging. Behaviorally and physiologically, these flies phenocopied naturally aged flies, supporting the notion that changes within the primary taste neurons underlie age-related loss of sugar taste in flies, rather than downstream processing or motor mechanisms. In agreement with these findings, aging in mammals directly impacts taste receptor cells [52].

We find that both natural aging and expression of Arctic in taste neurons lead to impaired sucrose response without affecting fatty acid taste. Sugars and fatty acids respond by different intercellular signaling pathways and receptors, with sugars activating Gsα [32] and fatty acids activating a complex set of intercellular signaling molecules including *norpA/Phospholipase C* (PLC; [34]). These findings raise the possibility that Gsα is more sensitive to the effects of aging, while PLC is more resilient. Consistent with this notion, aged mice display a reduction in the sugar receptor TAS1R, but not PLC [52]. While these findings raise the possibility that cAMP signaling is selectively reduced in *Gr64f* neurons, we did not observe significant expression differences in genes associated with sugar or fatty acid taste. The use of genetically encoded reporters, including the cAMP reporters cAMPr and ePAC, provide the opportunity to directly test the hypothesis that age selectively impacts signaling pathway(s) associated with sugar sensing [60,61].

A growing body of literature supports the notion that flies expressing human AD pathology-associated variants recapitulate many of the phenotypes associated with the disorder. For example, expression of Aβ variants, including Arctic, result in reductions in sleep [17,62], memory [63,64], and longevity [20,21,65]. Most studies to date have used broad expression of Aβ, either in all neurons or large neural structures within the brain. Therefore, little is known about the cell-autonomous impacts of Aβ expression on *Drosophila* behavior. Our finding that the expression of Arctic in sweet taste neurons impairs sugar taste response provides an additional application to study the cell type-specific effects of human Aβ variants on neuronal function and physiology. We find that selective expression of Arctic in taste neurons results in a decrease in SEZ innervation. We reported these results by monitoring mCD8GFP, whose axonal transport is mediated by an actin- and vesicle-based trafficking system that regulates delivery of synaptic components to target synapses [66]. This is consistent with our findings in which delivery of brp to synaptic terminals in the SEZ is impaired in flies expressing Arctic in *Gr64f* neurons. Therefore, it is surprising that these flies maintain their response to fatty acids. It is possible that either the subset of neurons that are sensitive to fatty acids are resilient to the effects of Arctic expression, or that only a subset of *Gr64f* neurons are required for fatty acid taste.

Here, we identify several candidate genes that are differentially expressed in *Gr64f+* neurons between young and aged flies. For example, multiple OBPs are downregulated in aging flies, and while these genes are broadly known to modulate olfaction, they are also expressed in the taste sensilla of insects [67]. While less is known about the regulation of taste by OBPs, one of the proposed functions of OPBs in the taste system is to modulate the clearance of bitter compounds. We find that *obp18a* is downregulated in *Gr64f+* neurons of aged flies. A previous study demonstrated that downregulation of *obp18a* is associated with decreased consumption of bitter tastants [68]. Although gustatory OPBs have been previously mapped to thecogen support cells [69], it is possible they serve diverse functions across multiple cell types. Alternativity, it is possible that OBPs interact directly with sweet taste neurons to modulate their activity, as is the case with *obp49*, which has been shown to be required for suppression of sweet tastants upon exposure to bitter substances [69]. Medium-chain fatty acids activate both bitter and sweet gustatory receptor neurons, raising the possibility that the age-dependent shift in OBP expression contributes to the decrease in sugar sensitivity with age [31]. Further study of how *obp18a* and other OBP family genes contribute to sugar taste during aging may inform our understanding of why there is a reduction in sugar, but not fatty acid taste.

We also identify numerous genes that are upregulated with aging. The finding that the immediate early gene *Arc1* is upregulated in *Gr64f+* neurons of aged flies is of particular interest. *Arc1* has complex functions in *Drosophila* including the mediation of transsynaptic RNA trafficking [44]. It is possible that either upregulation of *Arc1* reflects an overall reduction in *Gr64f* neuron activity, or that there is altered communication between gustatory receptor neurons and their targets in the SEZ. Together, the 66 genes that are differentially expressed provide the basis for future screening and analysis of age-dependent changes in taste neurons.

Taken together, we have found that aging is associated with reduced sensitivity to sugars but not fatty acids. These findings suggest that the intracellular signaling properties associated with sugar and fatty acid taste are differentially impacted during aging. While expression of an Aβ_1-42_ variant results in similar behavioral and physiological phenotypes as naturally aged flies, the effects on the connectivity of *Gr64f* neurons differs. Together, these findings establish the taste system as a model to investigate age- and neurodegenerative disease-associated reductions in sensory function.

## METHODS

### Fly husbandry and maintenance

Flies were grown and maintained on standard food media (Bloomington Recipe; Genesee Scientific, San Diego, CA). Flies were housed in incubators (Powers Scientific; Warminster, PA, USA) on a 12:12 LD cycle at 25°C with a humidity level of 55–65%. The following fly strains were ordered from the Bloomington Stock Center: *w*^1118^ (#5905; [70]), *Gr64f*-GAL4 (#57668; [14]), UAS-mCD8GFP (#32186; [71]), UAS-Aβ_1-40_ (#64215; [28]). UAS-Arctic was provided by Matt Kayser and was previously described in [21], UAS-GCaMP-R was provided by Greg Macleod and was previously described in [37], and UAS-Brp-Short was provided by Tim Mosca and was previously described in [42]. All flies were backcrossed to the *w*^1118^ genetic background for a minimum of 6 generations prior to testing. For aging experiments, *w*^1118^ flies were isolated 1 day post-eclosion, sorted by sex into vials of ∼30 flies, and then transferred to new vials every other day until they reached the indicated age for each experiment.

### Reagents

Fatty acids were obtained from Sigma Aldrich (St Louis, MO, USA): hexanoic acid (6C; #21530), octanoic acid (8C; #O3907). Sugars were purchased from Fisher Scientific (Hampton, New Hampshire, USA): sucrose (#FS S5-500), fructose (#57-48-7). All tastants were also dissolved in water.

### Proboscis Extension Response

Female flies were starved for 24 h prior to each experiment and then PER was measured as previously described [11,72]. In experiments using males, flies were starved for 18 h prior to each experiment. Briefly, flies were anesthetized with CO_2_ and then glued to a microscope slide (#12-550-15; Fisher Scientific) so that their head and proboscis were free to move while their thorax and abdomen remained restrained. After a 60 min acclimation period in a humidified box, flies were presented with water and allowed to drink freely until satiated. Flies that did not stop responding to water within 5 min were discarded. A wick made of Kimwipe (#06-666; Fisher Scientific) was placed partially inside a capillary tube (#1B120F-4; World Precision Instruments; Sarasota, FL) and then saturated with tastant. The saturated wick was then manually applied to the tip of the proboscis for 1–2 s and proboscis extension reflex was monitored. Only full extensions were counted as a positive response. Each tastant was presented a total of three times, with 1 min between each presentation. Both sugars and fatty acids were dissolved in water and tested at the indicated concentration. PER was calculated as the percentage of proboscis extensions divided by the total number of tastant presentations. For example, a fly that extends its proboscis twice out of the three presentations will have a PER response of 66%. Experiments were run ∼3 times per week until completion. Genotype and tastant presentation were randomized to ensure data reproducibility.

### Immunohistochemistry

For innervation and active zone measurements, the brains of female flies were dissected in ice cold phosphate buffered saline (PBS) (#BP243820; Fisher Scientific) and fixed in 4% formaldehyde, PBS, and 0.5% Triton-X for 30 min at room temperature. Brains were rinsed 3X with PBS and 0.5% Triton-X (PBST) for 10 min at room temperature and then mounted in Vectashield (VECTOR Laboratories; Burlingame, CA). For the representative image of Gr64f expression, brains were dissected in PBS and then fixed for 30 min at room temperature. Brains were rinsed 3X with PBST for 10 min then incubated overnight at 4°C. The next day, brains were incubated in primary antibody (1:20 mouse nc82; Iowa Hybridoma Bank; The Developmental Studies Hybridoma Bank; Iowa City, Iowa, USA) diluted in 0.5% PBST at 4°C for 48 hrs. Next, the brains were rinsed 3X with PBST for 10 min at room temperature and placed in secondary antibody (1:400 donkey anti-mouse Alexa Fluor 647; #A-31571; ThermoFisher Scientific; Waltham, Massachusetts, USA) for 90 min at room temperature. The brains were again rinsed 3X with PBST for 10 min at room temperature and then mounted in Vectashield. All brains were imaged in 1 μm z-plane interval sections on a Nikon A1R confocal microscope (Nikon; Tokyo, Japan) using a 20X oil immersion objective.

### Quantification of volume and fluorescence intensity

Quantification of mCD8GFP or Brp-short was performed by generating a sum intensity projection of the *Gr64f* neurons. A threshold of 491 (low) to 65535 (high) was applied to all images prior to quantification. One region of interest (ROI) was drawn around the dorsal taste peg and internal mouthpart projections while a second ROI was drawn around the ventral labellar projections. For volume measurements, the sum of the total number of fluorescent pixels in each slice was used. For intensity measurements, the mean of the intensity averaged from each slice was used. Images are presented as the Z-stack projection through the entire brain and processed using Fiji [73].

### In vivo calcium imaging

Flies expressing UAS-GCaMP-R (GCaMP6.0 and mCherry) in *Gr64f* neurons were starved for 24 h prior to imaging, as previously described [31]. Flies were anesthetized on ice and then restrained inside of a cut 200 μL pipette tip so that their head and proboscis were accessible, while their body and tarsi remained restrained. The proboscis was manually extended and then a small amount of dental glue (#595953WW; Ivoclar Vivadent Inc.; Amherst, NY) was applied between the labium and the side of the pipette tip, ensuring the same position throughout the experiment. Next, both antennae were removed. A small hole was cut into a 1 cm^2^ piece of aluminum foil and then fixed to the fly using dental glue, creating a sealed window of cuticle exposed. Artificial hemolymph (140mM NaCl, 2mM KCl, 4.5mM MgCl2, 1.5mM CaCl2, and 5mM HEPES-NaOH with pH = 7.1) was applied to the window and then the cuticle and connective tissue were dissected to expose the SEZ. Mounted flies were placed on a Nikon A1R confocal microscope and then imaged using a 20X water-dipping objective lens. The pinhole was opened to allow a thicker optical section to be monitored. *Gr64f* neurons were simultaneously excited with wavelengths of 488 nm (FITC) and 561 nm (TRITC). All recordings were taken at 4 Hz with 256 resolution. Like PER, tastants were applied to the proboscis for 1–2 s with a wick, which was operated using a micromanipulator (Narishige International USA, Inc.; Amityville, NY). Experiments were run ∼3 times per week until completion.

### Quantification of calcium activity

To quantify UAS-GCaMP-R activity, regions of interest were first drawn manually around the *Gr64f* projections of one hemisphere, alternating between flies. For each frame, the mean fluorescence intensity for FITC and TRITC was subtracted from background mean fluorescence intensity. Then, the fluorescence ratio of GCaMP6.0 to mCherry was calculated. Next, baseline fluorescence was calculated as the average fluorescence ratio of the first 5 frames, 10 s prior to tastant application. For each frame, the % change in fluorescence (%ΔF/F) was then calculated as: [(peak fluorescence ratio -baseline fluorescence ratio) / baseline fluorescence ratio] * 100. Average fluorescence traces were created by taking the average and standard error of %ΔF/F for each recording of a specific tastant.

### Single-nucleus RNA sequencing

#### Tissue collection, nuclei isolation, library preparation, and sequencing

Fly labellums were dissected at 10 and 40 days and then placed into 1.5 mL RNase-free Eppendorf tubes, flash-frozen in liquid nitrogen, and then stored at -80°C. Approximately ∼400 labellum were collected for each treatment and stored on dry ice. Nuclei isolation and snRNAseq were performed by Singulomics Corporation (Singulomics.com; New York). To isolate nuclei, tissue was homogenized and lysed with Triton X-100 in RNase-free water. The nuclei were then purified, centrifuged, resuspended in PBS with RNase Inhibitor, and diluted to 700 nuclei/μL. Standardized 10x capture and library preparation were performed using the 10x Genomics Chromium Next GEM 3’ Single Cell Reagent kit v3.1 (10x Genomics; Pleasanton, CA). The libraries were then sequenced with Illumina NovaSeq 6000 (Illumina; San Diego, CA). The snRNA-seq raw sequencing files were processed with CellRanger 6.0 (10x Genomics; Pleasanton, CA). The sequencing reads were mapped to the *Drosophila* reference genome (Flybase r6.54).

#### Data normalization and integration

All analysis of snRNA-seq data was performed in R (v4.3) using Seurat (v4.3.0; [74]). Unless otherwise noted, default parameters were used for all operations. Each sample was independently normalized using the SCTransform function (v2;) [75,76]. Following normalization, integration features were selected using the SelectIntegrationFeatures function, with nFeatures = 3000. Normalized data was prepared for integration with the PrepSCTIntegration function, using the features identified in the previous step. Integration anchors were identified using the FindIntegrationAnchors function, with normalization.method = “SCT”. Finally, the data was integrated with the IntegrateData command, using the previously identified anchors.

#### Cell type clustering and dimensionality reduction

Following integration, Principal Component Analysis was performed using the RunPCA function, with npcs = 30. The resulting PCs were used to identify clusters, with a resolution of 0.3, resulting in 13 total clusters. The integrated data was prepared for marker identification by running the PrepSCTFindMarkers function, and then markers were identified with the FindAllMarkers command, using the MAST statistical test [77], which has been found to perform well on single-cell data sets [78].

The list of marker genes was used to annotate cluster identity. First, we compared the marker genes for each cluster to previously generated datasets in *Drosophila*, using the Cell Marker Enrichment tool (DRscDB; https://www.flyrnai.org/tools/single_cell/web/enrichment[79]). Wherever possible, we refined our labels by comparing marker genes with published literature marking known cell types [80].

#### Differential gene expression (DEG) aging analysis

To identify genes in *Gr64f*+ nuclei that are differentially expressed in aged flies, we applied the FindMarkers function on these cells, again using the MAST statistical test, and set group.by=‘age’. Genes with an adjusted *P* value less than 0.05 and an average Log_2_ Fold Change greater than 0.25 or less than -0.25 were considered differentially expressed and used for downstream analysis.

#### Gene Ontology (GO) analysis

Gene ontology pathway analyses were conducted in R (v.4.3.0), using the clusterProfiler package (v4.8.3; [81]). For gene set enrichment analyses, a log_2_FC value was calculated between the young (10d) and aged (40d) groups for each gene expressed in the cell population of interest. The sorted log_2_FC list was passed to the gseGO function (with options: ont = ‘BP’, keyType = ‘SYMBOL’, minGSSize = 3, maxGSSize = 800, OrgDb = org.Dm.eg.db, pvalueCutoff = 0.01). The results of this test were passed to the dotplot function for visualization.

### Statistical Analysis

All measurements are presented as bar graphs or line graphs showing mean ± standard error. Measurements of PER were not normally distributed; therefore, the non-parametric restricted maximum likelihood (REML) estimation was used. For all other measurements, either a t-test or a two-way analysis of variance (ANOVA) was used. Post hoc analyses were performed using Sidak’s multiple comparisons test. Statistical analyses and data presentation were performed using InStat software (GraphPad Software 8.0; San Diego, CA). Sample sizes for behavioral and functional imaging experiments are consistent with previous studies [31]. Generally, ∼30–50 flies were used for each experimental or control group for behavioral experiments and ∼10 flies per group for imaging experiments.

## Acknowledgments

We would like to thank members of the Keene and Dahanukar labs for technical assistance and helpful discussion. This work was supported by the National Institutes of Health (NIH) grants R01NS085252 to ACK, R01DC017390 to ACK and AD, as well as R00AG071833 to EBB. This work was also supported by the NIH T32 grant DC000044 and the National Science Foundation (NSF) REU grant DBI-1852175.

## FIGURE LEGENDS

## SUPPLEMENTARY FIGURE LEGENDS

**Figure S1. Taste response to additional tastants in aging and AD model flies. (A)** There is a significant effect of age on PER to fructose in *w*^1118^ flies (two-way ANOVA: F_1,396_ = 11.82, *P* < 0.0006; N = 28–40). **(B)** There is a significant effect of Arctic expression on PER to fructose (two-way ANOVA: F_2,726_ = 21.54, *P* < 0.0001; N = 34–50). **(C)** There is no effect of Aβ_1-40_ expression on PER to sucrose (two-way ANOVA: F_2,858_ = 2.779, *P* < 0.1232; N = 47–50). **(D)** There is no effect of age on PER to octanoic acid in *w*^1118^ flies (two-way ANOVA: F_1,468_ = 0.1367, *P* < 0.7117; N = 40). **(E)** There is no effect of Arctic expression on PER to octanoic acid (two-way ANOVA: F_2,582_ = 0.2824, *P* < 0.7541; N = 20–40). **(F)** There is no effect of Aβ_1-40_ expression on PER to hexanoic acid (two-way ANOVA: F_2,800_ = 0.1829, *P* < 0.8329; N = 37–50). Error bars indicate ± SEM. * *P* < 0.05; ** *P* < 0.01; *** *P* < 0.001.

**Figure S2. Taste response to sugar, but not fatty acids, is reduced at intermediate concentrations of sucrose and hexanoic acid in aging and AD model male flies. (A)** There is a significant effect of age on PER to sucrose in *w*^1118^ flies (two-way ANOVA: F_1,472_ = 30.77, *P* < 0.0001; N = 34–40). **(B)** There is no effect of age on PER to hexanoic acid in *w*^1118^ flies (two-way ANOVA: F_1,408_ = 0.6961, *P* < 0.4046; N = 30–40). **(C)** There is a significant effect of Arctic expression on PER to sucrose (two-way ANOVA: F_2,678_ = 21.98, *P* < 0.0001; N = 36–40). **(D)** There is no effect of Arctic expression on PER to hexanoic acid (two-way ANOVA: F_2,700_ = 0.3514, *P* < 0.7039; N = 38–40). Error bars indicate ± SEM. * *P* < 0.05; ** *P* < 0.01; *** *P* < 0.001; **** *P* < 0.0001.

**Figure S3. Quality assessment of snRNA-seq results**. The **(A)** total number of Unique Molecular Identifiers (UMI), **(B)** number of expressed genes, and **(C)** percentage of mitochondrial transcripts are shown for each replicate (rep).

**Figure S4. snRNA-seq clustering and differential gene expression analysis. (A)** Dot plot of the top 3 marker genes in each cluster based on log fold-change in expression. **(B)** Volcano plot depicting differentially expressed genes within the sensory cluster between 10- and 40-day-old flies. **(C)** Gene Ontology (GO) analysis of the differentially expressed genes identified in **(C)**.

## SUPPLEMENTARY TABLE LEGENDS

**Supplementary Table 1. Marker genes and annotations by cluster**. For each gene, the *P* value, Log_2_ (Fold Change), Adjusted *P* value, and indicated cluster are listed. Also listed is the percentage of nuclei within each cluster that express a given gene as well as the percentage of nuclei outside the cluster.

**Supplementary Table 2. Differentially expressed genes within the sensory neu bron cluster between young (10d) and aged (40d) samples**. For each gene, the *P* value, Log_2_ (Fold Change), and Adjusted *P* value are listed. Also listed is the percentage of nuclei within the sensory neuron cluster that express a given gene as well as the percentage of nuclei outside the sensory neuron cluster.

**Supplementary Table 3. Gene ontology analysis of differentially expressed genes within the sensory neuron cluster between young (10d) and aged (40d) samples**. (Description needs to be added)

**Supplementary Table 4. Differentially expressed genes in *Gr64f*+ nuclei between young (10d) and aged (40d) samples**. For each gene, the *P* value, Log_2_ (Fold Change), and Adjusted *P* value are listed. Also listed is the percentage of nuclei within *Gr64f*+ nuclei that express a given gene as well as the percentage of nuclei that do not express *Gr64f*.

**Supplementary Table 5. Gene ontology analysis of differentially expressed genes within the sensory neuron cluster between young (10d) and aged (40d) samples**. (Description needs to be added)

