## Supplementary figures and images for "Aging is associated with a modality-specific decline in taste"

### Supplemental Figure 1

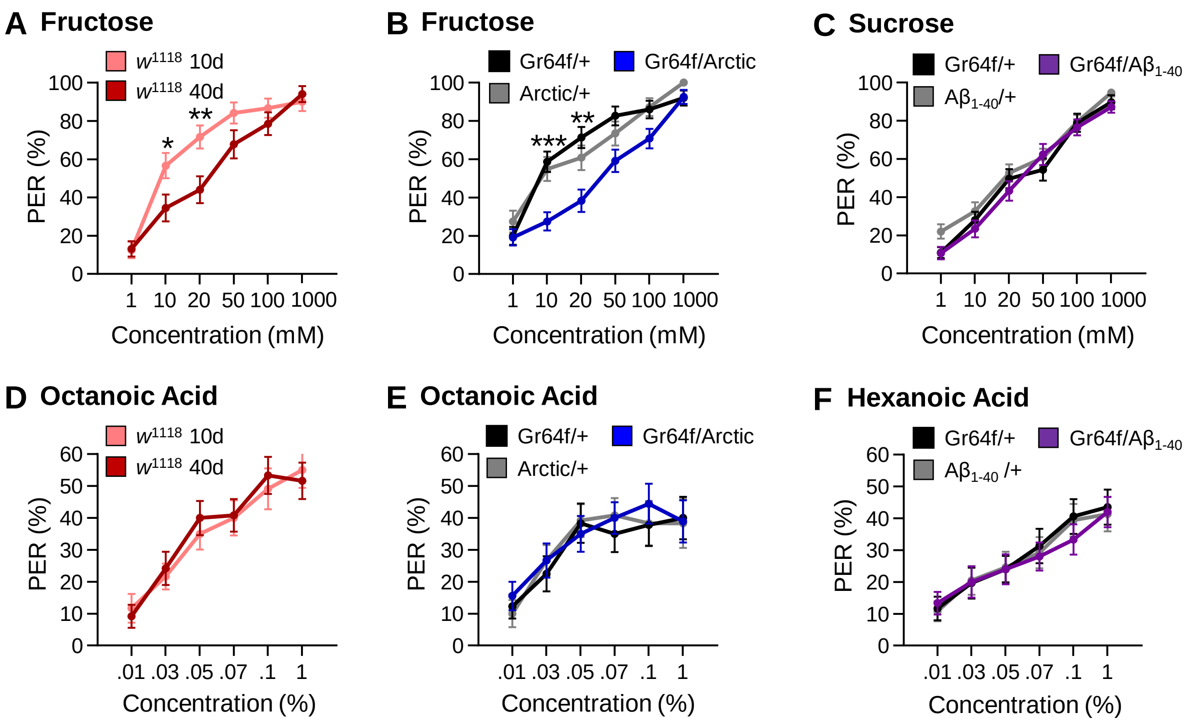

### Supplemental Figure 2

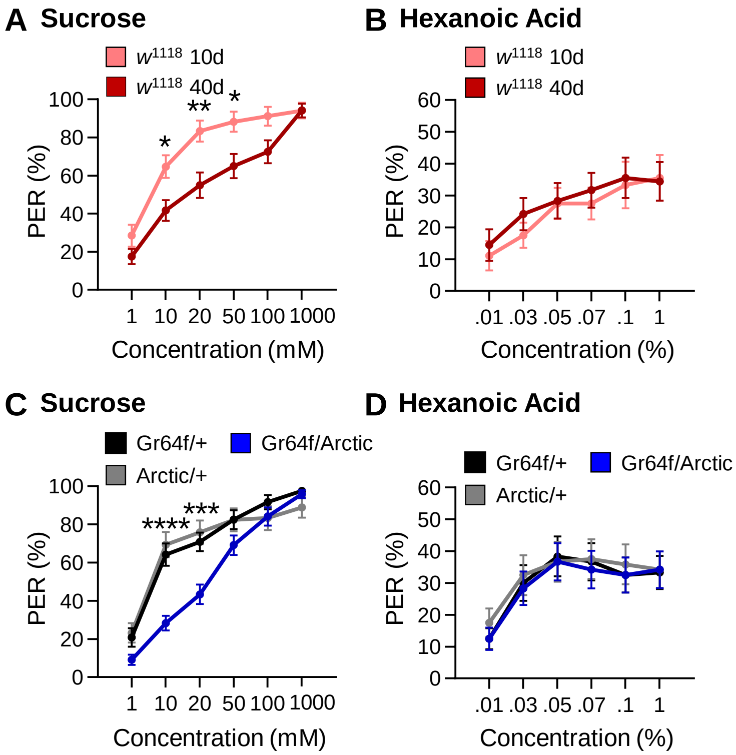

### Supplemental Figure 3

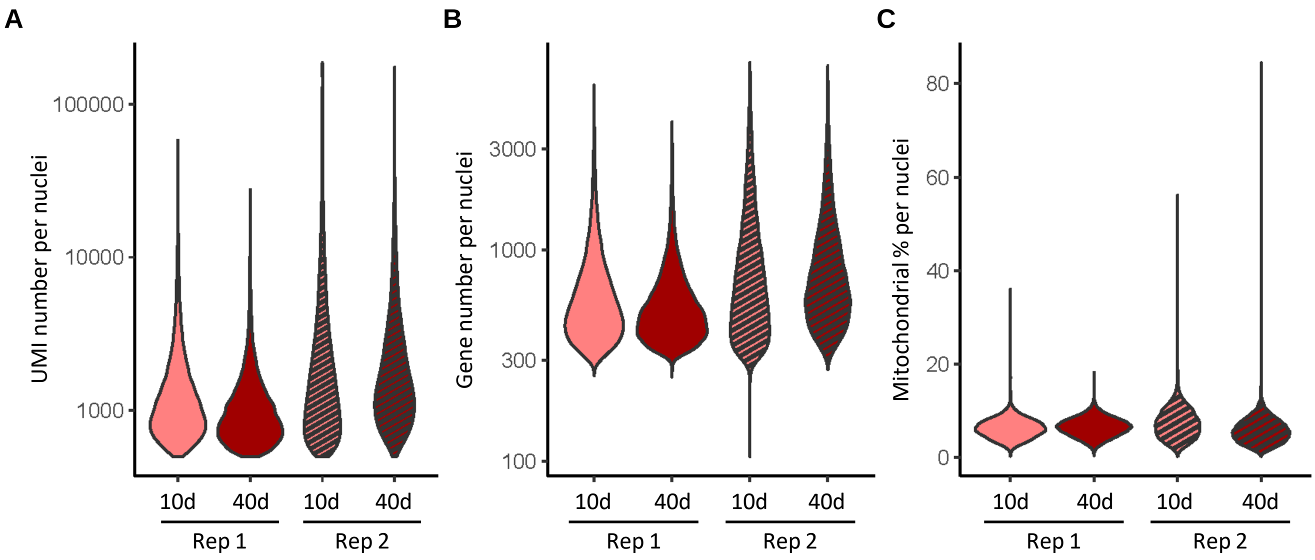

### Supplemental Figure 4

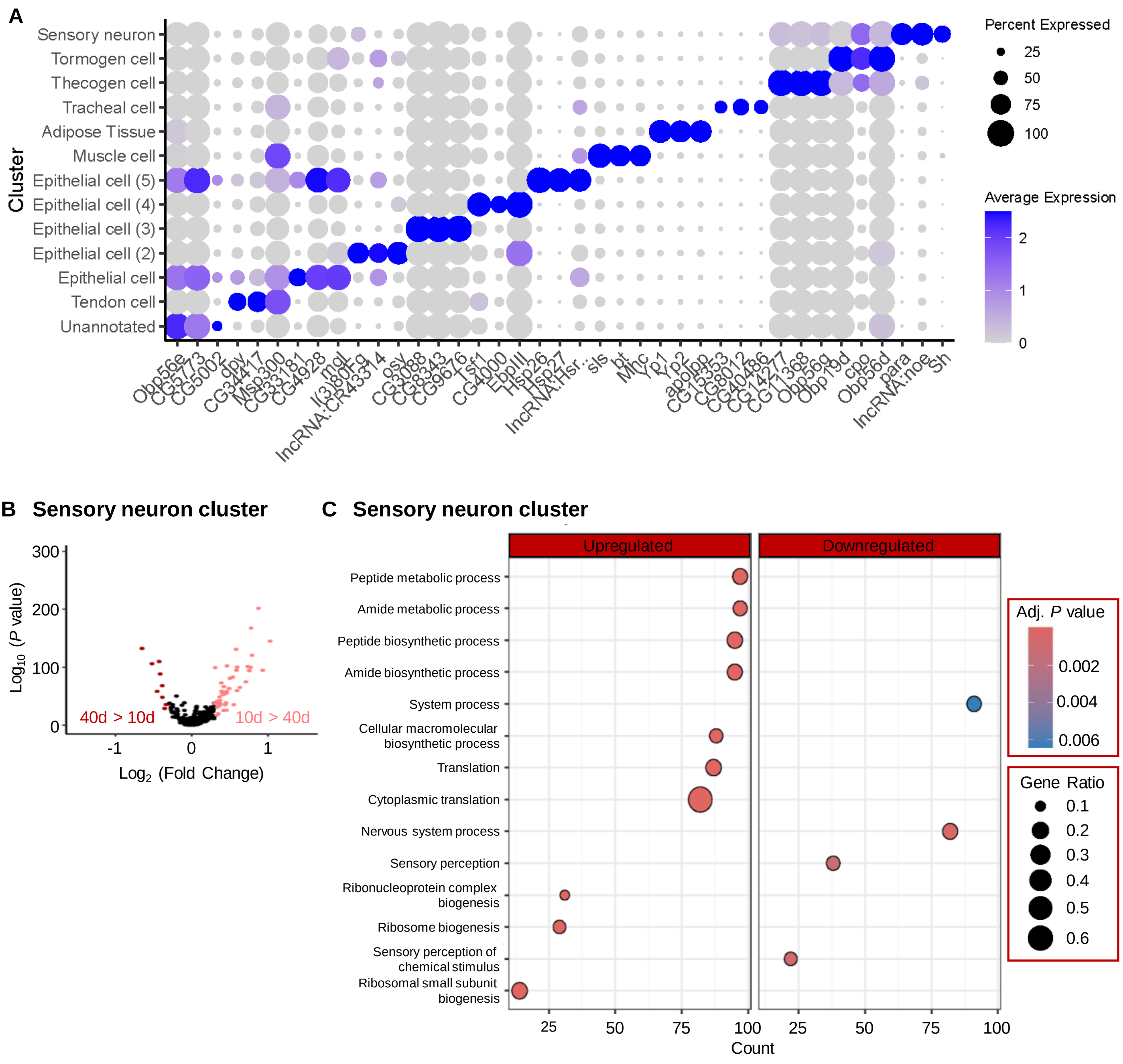
